# Cutin-Derived Oligomers Act as Damage-Associated Molecular Patterns in *Arabidopsis thaliana*

**DOI:** 10.1101/2023.05.16.540997

**Authors:** Carlos J.S. Moreira, Rita Escórcio, Artur Bento, Marta Bjornson, Ana S. Tomé, Celso Martins, Mathieu Fanuel, Isabel Martins, Benedicte Bakan, Cyril Zipfel, Cristina Silva Pereira

## Abstract

The cuticle constitutes the outermost defensive barrier of most land plants. It comprises a polymeric matrix – cutin, surrounded by soluble waxes. Moreover, the cuticle constitutes the first line of defense against pathogen invasion, while also protecting the plant from many abiotic stresses. Aliphatic monomers in cutin have been suggested to act as immune elicitors in plants. This study analyses the potential of tomato cutin oligomers to act as damage-associated molecular patterns (DAMPs) able to induce a rapid immune response in the model plant *Arabidopsis*. Cutin oligomeric mixtures led to Ca^2+^ influx and MAPK activation in *Arabidopsis*. Comparable responses were measured for cutin, which was also able to induce a reactive oxygen species (ROS) burst. Furthermore, treatment of *Arabidopsis* with cutin oligomers resulted in a unique transcriptional reprogramming profile, having many archetypal features of pattern-triggered immunity (PTI). Targeted spectroscopic and spectrometric analyses of the cutin oligomers suggest that the elicitors compounds consist mostly of two up to three 10,16-dihydroxyhexadecanoic acid monomers linked together through ester bonds. This study demonstrates that cutin breakdown products can act as DAMPs; a novel class of elicitors deserving further characterization.

## Introduction

Plants occupied land environments approximately 450 million years ago (Delwiche and Cooper, 2015). The transition from water to land habitats exposed plants to numerous challenges imposed by an extremely desiccating environment (Waters, 2003). To control water loss, protect against UV radiation and pathogens, and reinforce the epidermal cell layer, plants developed a hydrophobic barrier – the cuticle (Martin and Rose, 2014; Fich et al., 2016). The cuticle is composed of a polymeric matrix of cutin to which organic solvent soluble lipids (waxes) associate (Yeats and Rose, 2013). In addition, cutin interaction with the polysaccharides that build up the epidermal cell walls has been proposed (Philippe et al., 2020a; Xin and Fry, 2021), but the nature of such anchoring remains uncertain.

During infection of the aerial organs of plants, fungal spores release cutin-degrading enzymes – cutinases, having esterase activity (Longhi and Cambillau, 1999) that can disrupt the polymeric matrix and release cutin-derived molecules. Perception of these molecules by the fungus increases the production of cutinases that breach the cuticle barrier, thus allowing the fungus to invade the plant organ (Kolattukudy et al., 1995).

To fend off pathogen invasion, plants have developed a highly specialized mechanism to sense biotic threats by using cell surface pattern recognition receptors (PRRs)(Zipfel, 2014). These receptors perceive pathogen-associated molecular patterns (PAMPs) and damage-associated molecular patterns (DAMPs), derived from the invading pathogens or from the breakdown of plant tissues, respectively (Zipfel, 2014). Cutin aliphatic monomers (*i.e.* the major basic elements composing the cutin polymer) have been proposed as DAMPs due to their ability to induce some elements of a canonical immune response, namely the production of reactive oxygen species (ROS) in cucumber, rice and *Arabidopsis thaliana* (hereafter *Arabidopsis*), and the upregulation of defense-related genes in rice and *Arabidopsis* (Kauss et al., 1999; Kim et al., 2008; Park et al., 2008). Exogenous application of monomers obtained from plants having augmented cuticular permeability (*SlSHN3*-OE), increased the resistance of Micro-Tom tomato plants against the fungal pathogen *Botrytis cinerea*, and activated defense responsive genes (Buxdorf et al., 2014); however, the nature of the elicitor(s) remains unresolved. Cutin aliphatic monomers were also reported to induce the production of antimicrobial compounds (Serrano et al., 2014). Although cutin monomers have been proposed as DAMPs, their capabilities to elicit other important hallmark early immune responses, for example intracellular calcium influx and activation of mitogen-activated protein kinases (MAPKs), have never been observed (Serrano et al., 2014; Hou et al., 2019). Also, it is unclear if the tested cutin aliphatic monomers are the most potent class of cutin-derived DAMPs. Esterase-based degradation of cutin progresses through ester-cleavage, likely releasing cutin oligomers and not only monomers (Beneloujaephajri et al., 2013). This raises the hypothesis that cutin oligomers act as DAMPs, similar to that proposed for cutin monomers.

To investigate the hypothesis that cutin oligomers (COMs) can activate PTI responses, the cutin polymer was first isolated from tomato peel (Moreira et al., 2020; Bento et al., 2021), and subsequently broken down through a mild chemical hydrolysis to generate COMs. The ability of the produced COMs to activate calcium influx, MAPK activation and transcriptional reprograming in *Arabidopsis* was investigated. The results clearly indicate that COMs act as DAMPs. Spectroscopy and spectrometry analyses suggested that the elicitors are dimers and/or trimers consisting mostly of esterified 10,16- dihydroxyhexadecanoic acid units (dihydroxy-C16 acid), one of which possibly methylated. The hypothesis that cutin disruption releases oligomers able to act as DAMPs is discussed in detail.

## Results and discussion

### Cutin polymer activates a ROS burst in Arabidopsis

We hypothesized that the degradation of the plant polyester cutin is coordinated with the release of polymeric/oligomeric variants capable of eliciting hallmark early plant immune responses. We first tested a cutin polymeric variant ability to induce a ROS burst in *Arabidopsis*. *Arabidopsis* is a well-established model plant for studying PTI due to the diversity of established protocols and plant resources, including characterized mutants and reporter lines (Felix et al., 1999). The ROS burst was measured through a well-established protocol that detects a luminescence signal produced in the presence of H_2_O_2_ due to peroxidase (here, horseradish peroxidase, HRP)-mediated conversion of luminol (Zhu et al., 2016).

To obtain the cutin, an ionic liquid extractant was applied to isolate a highly pure cutin polymer (hereafter simply referred as cutin), showing minor ester cleavage (Moreira et al., 2020). This method ensures faster and simpler recovery of cutin compared to the conventional enzyme-based isolation (Moreira et al., 2020). Cutin was purified from tomato pomace since its high availability as an agroindustry residue (European Commission, 2021) enables the production of large amounts of polymeric structures. Moreover, previous studies showed that cutin purified with an ionic liquid from tomato pomace (consisting of peels, seeds and stems) is virtually similar to that obtained from the tomato peel fraction alone (Escórcio et al., 2022).

Exposure of seedlings of *Arabidopsis* to cutin (suspension in MiliQ water) resulted in a clear ROS burst (Fig. 1A-B and Supplemental Fig. S1). Flg22, a 22-amino acid peptide derived from bacterial flagellin, is a well-established strong inducer of plant immunity (Felix et al., 1999; Correia et al., 2020) (used here as positive control). The effect was reproducible, and the response was not depleted at 45 min post-treatment (Fig. 1B). On the contrary, pure compounds (commercially available), which are representative of tomato cutin constituents: long chain fatty acids, hydroxycinnamic acids or fatty acid monoglycerides having variable side chains, did not induce a ROS burst under the tested conditions (Fig. 1C, Supplemental Fig. S2). Cutin hydrolysates, which are obtained by an extensive hydrolysis of the cutin, consist almost exclusively of aliphatic monomers with a few aromatic monomers (Escórcio et al., 2022). These hydrolysates also did not elicit a ROS burst (Fig. 1D). Collectively, the results suggest that once the polymeric backbone of the plant polyester is deconstructed to its composing monomeric pieces, its capability to elicit a ROS burst is lost; hence some preservation of the polymeric backbone might be required for the eliciting of a ROS burst in *Arabidopsis*.

**Fig. 1.**
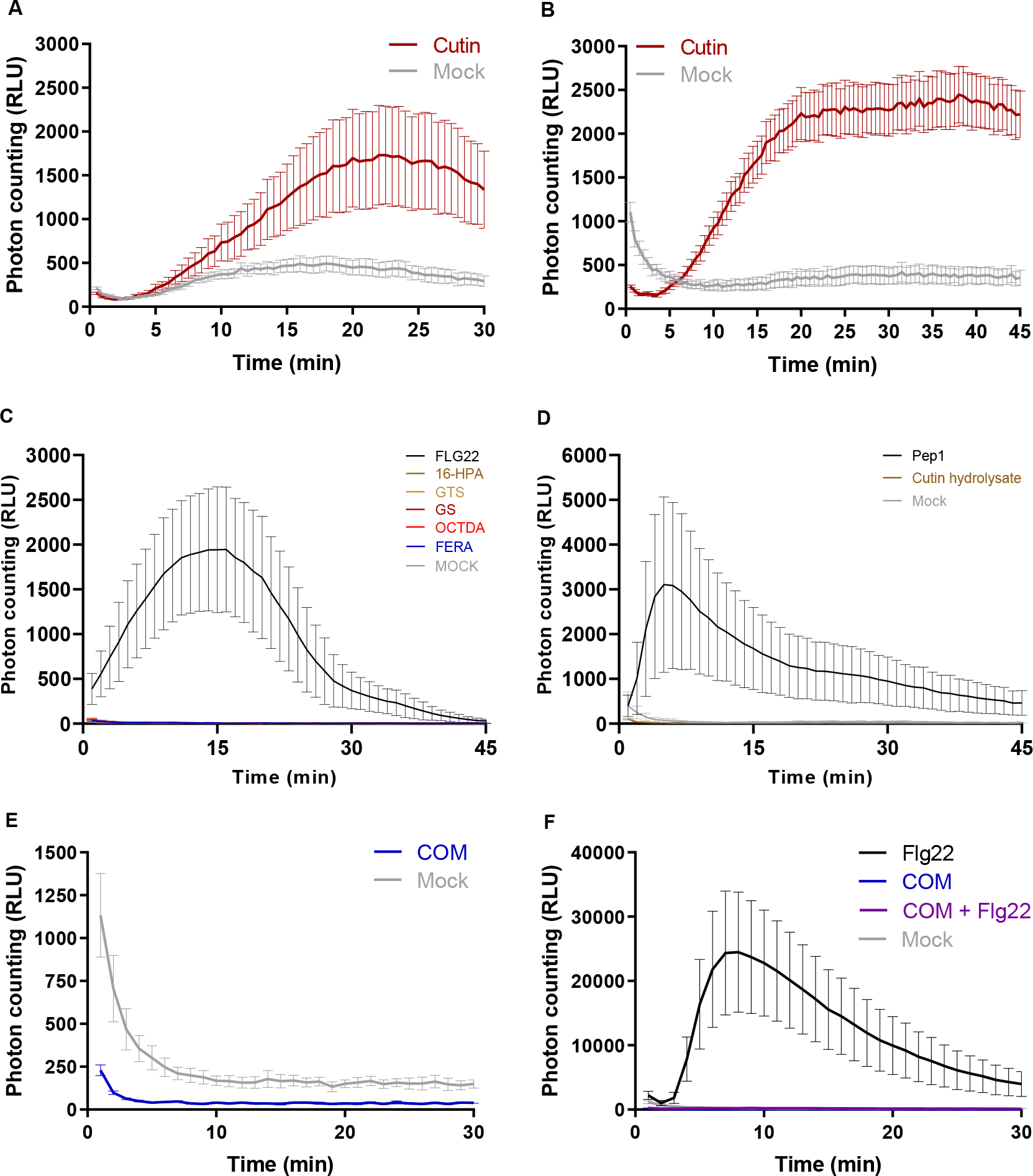
Luminescence-based detection of apoplastic ROS in *Arabidopsis thaliana* Col-0 leaf discs upon treatment with 2 mg·mL^-1^ of cutin in MiliQ water for 30 min (**A**) and 45 min (**B**); **C** – 1 mM of commercially available pure monomers (16- hydroxypalmitic acid (16-HPA), octanedioic acid (OCTDA) and ferulic acid (FERA)), and oligomers (glyceryl stearate (GS) and glyceryl tristearate (GTS)), in 10 % ethanol in MiliQ water for 45 min; **D** – 1 mg·ml^-1^ of cutin hydrolysate obtained after alkaline hydrolysis of cutin; **E** – 2 mg·ml^-1^ of COM obtained through the methanolysis of cutin in 0.5 % DMSO in MiliQ water; and **F** – co-treatment with 2 mg·ml^-1^ of COM and 100 nM Flg22 in 0.5 % DMSO in MiliQ water. In all the assays the Mock consists of the solvent. The positive controls were Flg22 (100 nM) or Pep1 (100 nM).

To test if small chains of monomers linked together through ester linkages; *i.e*. oligomers (<7), could act as elicitors, we prepared cutin oligomeric mixtures (COMs). To produce these, cutin was depolymerized through a mild chemical hydrolysis and the released molecules were collected (*see Materials and Methods*). The produced COMs were unable to trigger a ROS burst (Fig. 1E). However, no effect was detected when the seedings were co-treated with flg22 and COM, although flg22 alone clearly induced a ROS burst (Fig. 1F). This result suggests that constituents of the COM preparation interfered with the reporter of luminescence. In fact, the COM contains phenolic compounds (Supplemental, Table S1) and phenol oxidation has been reported to inactivate HRP activity in a concentration dependent mode (*i.e.* ratio enzyme:inhibitor) (Mao et al., 2013). Cutin hydrolysates (mixture of all hydrolysable cutin monomers) also contain low levels of phenolic compounds (Escórcio et al., 2022). There are alternative methods for ROS measurement; for example, DAB staining has been used to detect the accumulation of intracellular ROS in response to treatment with cutin aliphatic monomers, specifically hydroxy palmitic acid (HPA) (Kim et al., 2008; Park et al., 2008). Thus, while we observed that cutin aliphatic monomers did not induce an apoplastic ROS burst using a luminol-based assay, we cannot disregard the possibility that accumulation of intracellular ROS might occur at extended post-treatment periods.

The results show that treatment of *Arabidopsis* with cutin induced a ROS burst (Fig. 1A) but not any of the tested pure cutin constituents (Fig. 1C). Due to the aforementioned technical limitation, to further test COMs activity as potential elicitors of plant immunity, we converged towards the calcium response - another hallmark early immune response.

### Cutin and COMs, but not cutin hydrolysates, activate a calcium influx in Arabidopsis

Both cutin and COMs showed a clear and reproducible induction of calcium influx in *Arabidopsis* plants expressing aequorin (Fig. 2A, 2B), a widely used calcium activated reporter of immune responses in plants (Mithöfer and Mazars, 2002). The response patterns were however different: cutin response was bimodal with maximum values at 3 min and 12 min (Fig. 2A), whereas COM response was monomodal with a maximum between 3 and 5 min (Fig. 2B). This result suggests that cutin may comprise several classes of chemical elicitors, one of which is prevalent in the COM fraction. *Nicotiana benthamiana* plants expressing aequorin (Segonzac et al., 2011) when exposed to COMs also showed a calcium influx having a monomodal response-type (Supplemental Fig. S3A). Since the eliciting molecules were similarly recognized by both tested plants, the elicitors are likely not species-specific.

**Fig. 2.**
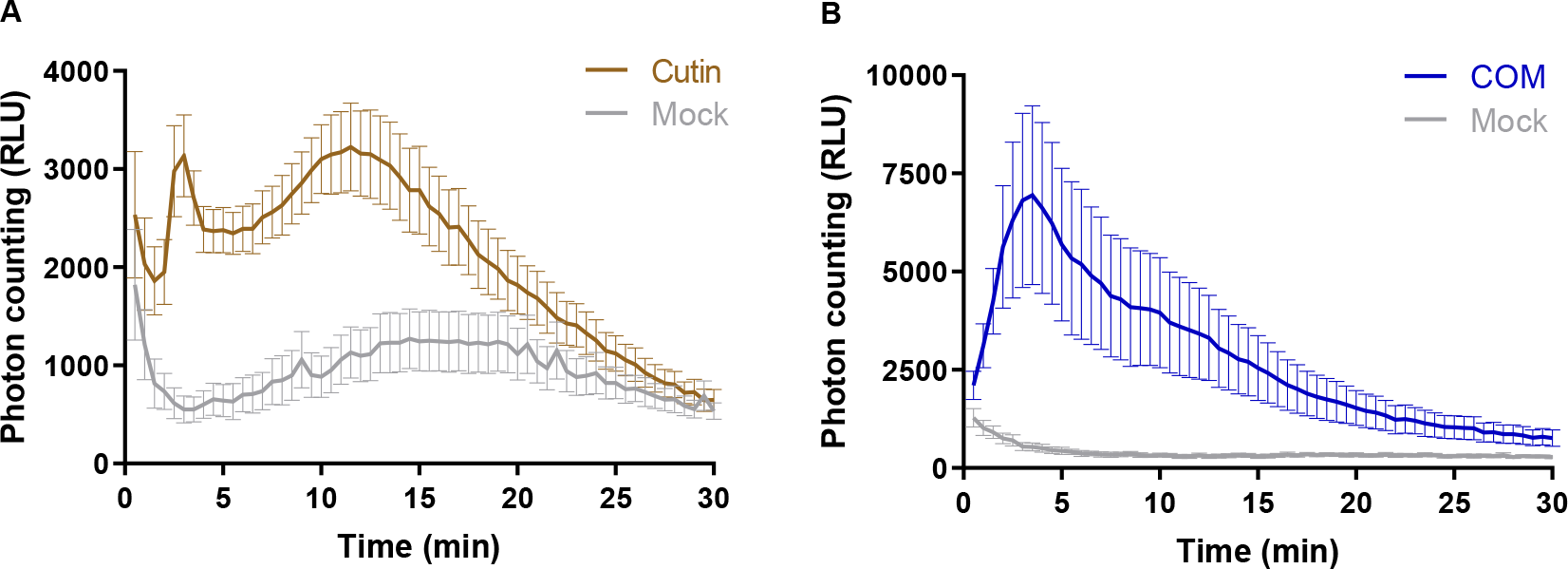
Luminescence-based detection of calcium influx in *Arabidopsis thaliana* seedlings expressing the calcium reporter aequorin, upon treatment with: **A** – 1 mg·mL^1^ of cutin in MiliQ water; **B** – 2 mg·mL^-1^ of COM in 0.5 % DMSO in MiliQ water. In all the assays the Mock consists of the solvent.

Finally, no induction of a calcium influx was observed upon treatment of *Arabidopsis* seedlings with either HPA or cutin hydrolysates increasing concentrations (Supplemental Fig. S3B-C). Collectively, these data validate the opening hypothesis that cutin small oligomers may act as elicitors of PTI. Mild deconstruction of the polymer potentiates its capability to induce a calcium influx in *Arabidopsis,* but its complete depolymerization abolished this eliciting effect.

### Cutin oligomers trigger MAPK activation in Arabidopsis

PTI signaling events occurring downstream to elicitor perception involve the activation of MAP kinases (Yu et al., 2017; DeFalco and Zipfel, 2021). Accordingly, the capabilities of COM to activate MAP kinases in *Arabidopsis* Col-0 seedlings was evaluated. This immunoblot-based assay allows the detection of the phosphorylated (active) forms of MAPK 3, 4, 6 and 11 during PTI signaling (Willmann et al., 2014). Short (10 min) exposure of *Arabidopsis* seedlings to COM activated hallmark MAPK activation; similar to that observed when the plants were exposed to flg22 (Fig. 3). In addition to the wild type plants, three mutants were also tested, single: *cerk1-2* (Ranf et al., 2011), double: *bak1-5 bkk1* (Roux et al., 2011), and triple: *bak1-5 bbk1 cerk1* (*bbc*) (Xin et al., 2016). CHITIN ELICITOR RECEPTOR KINASE 1 (CERK1) is a common co-receptor for LysM-type PRRs (Macho and Zipfel, 2014) while BRASSINOSTEROID INSENSITIVE 1-ASSOCIATED KINASE 1 (BAK1) and BAK1-LIKE KINASE 1 (BKK1) are common co-receptors for leucine-rich repeat- type PRRs (Tang et al., 2017). Thus, differences in the response pattern of the selected mutants may reveal potential families of PRR(s) that recognize the elicitor(s) within COM. The results showed that COM induced a clear activation of MAP kinases in all the mutants tested, similar to that observed in the wild-type plants (Fig. 3). The observation that MAPK activation was similar in all mutants is suggestive of a perception mechanism independent on the families of PRRs known to associate with CERK1 and SERKs. Ultimately, these results suggest that COMs triggered a MAPK-mediated signaling cascade, opening the hypothesis that COM exposure also involves transcriptional reprogramming.

**Fig. 3.**
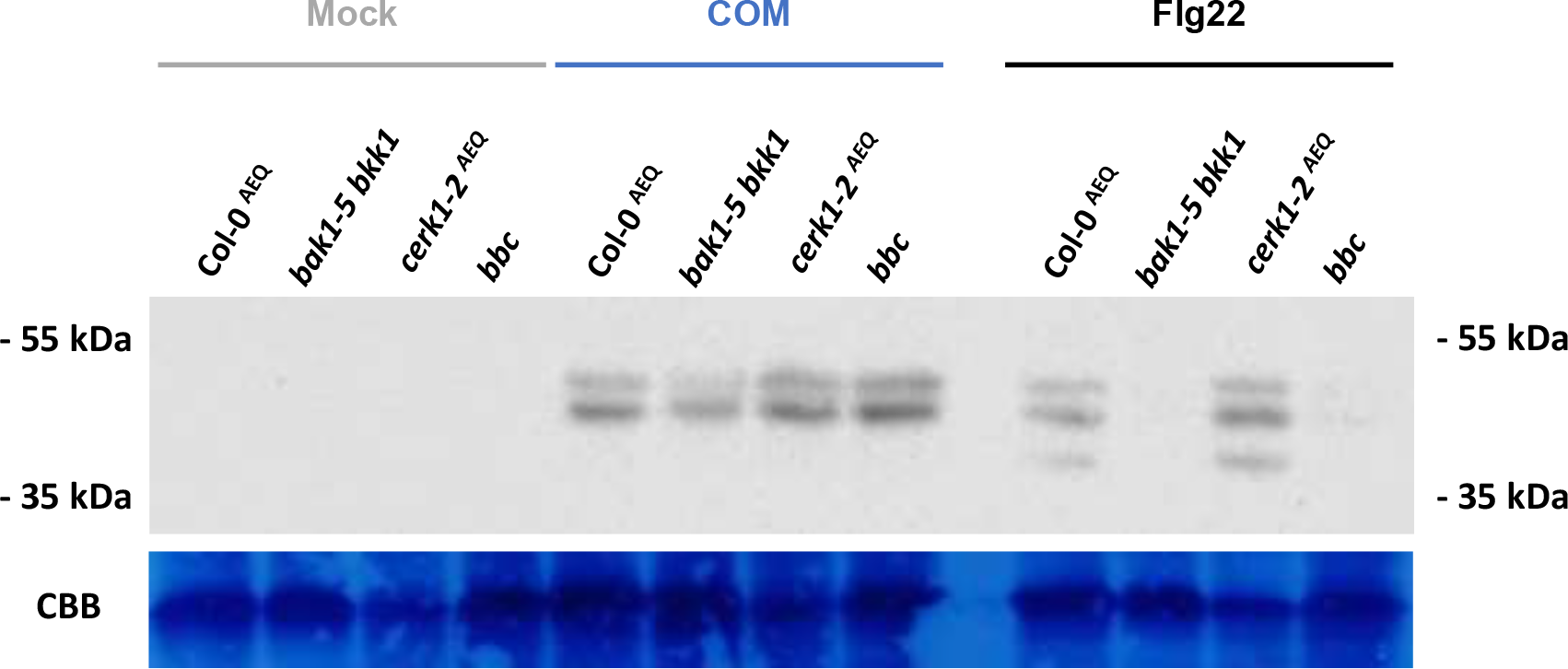
Western blot evaluation of MAPK activation in 14-day-old *Arabidopsis thaliana* seedlings from wild type Col-0^aeq^ plants and the *bak1-5 bkk1*, *cerk1-2^aeq^ and bbc* mutants, upon treatment with 3 mg·mL^-1^ of COM in 0.5 % DMSO in MiliQ water. The mock consists of the solvent, and the positive control was Flg22 (100 nM, in MiliQ water).

### Cutin oligomers treatment induced a transcriptional reprograming consistent with activation of PTI

We evaluated the transcriptional reprogramming in *Arabidopsis* seedlings upon a 30-min treatment with COM compared to mock control (RNA-seq). Previous studies covering distinct PTI elicitors showed significant responses at 30 min post-treatment (Bjornson et al., 2021). Principal components analysis (PCA) demonstrates that their transcriptomic profiles are clearly separating from each other (Supplemental Fig. S4). A total of 528 differentially expressed genes (DEGs) resulted from the COM treatment, of which 479 genes were upregulated, while only 49 were downregulated compared to the mock treatment (Fig. 4A). Enriched gene ontology (GO) categories were only obtained for the subset of upregulated genes due to the small number of downregulated genes. An enrichment for terms related to activation of plant immunity, particularly ‘response to wounding’ and ‘response to other organism’ was noticed (Fig. 4B).

**Fig. 4.**
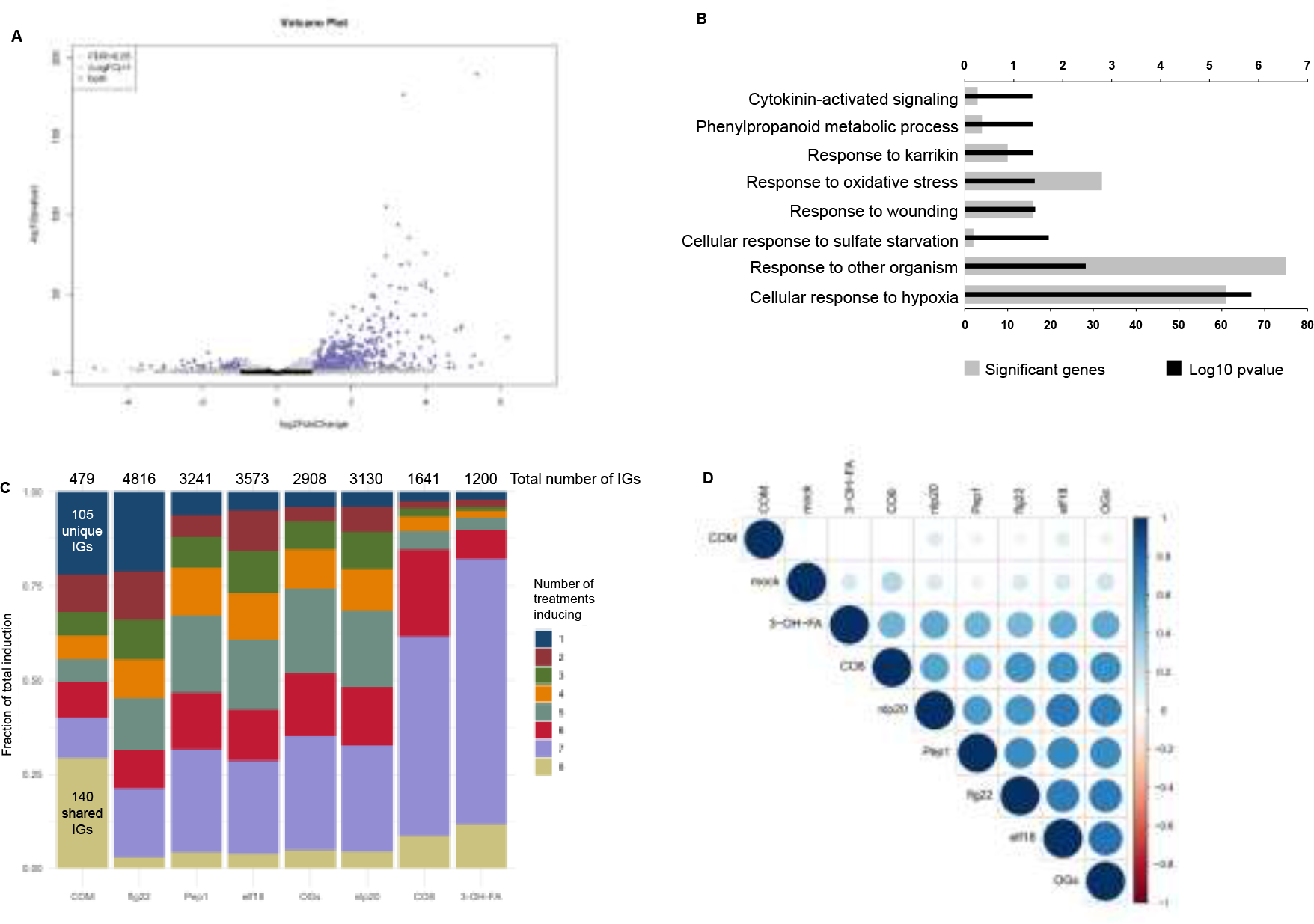
(A) Volcano plot representing the statistically significant (adjusted *p-*value < 0.05) differentially expressed genes (Log2 fold change ≤ -1 or ≥ 1), in *Arabidopsis thaliana* Col-0 plants upon treatment with COM for 30 min (479 upregulated genes and 49 downregulated genes). **(B)** GO term enrichment analysis of the genes that showed upregulation upon treatment with COM for 30 min. **(C)** Comparison of genes induced by treatment of COM with those induced by seven other elicitors of plant triggered immunity. The total number of induced genes (IGs) for each elicitor (over all time points in Bjornson, *et al*. 2021) is presented and the number of genes induced by all treatments or solely by COM are highlighted. **(D)** Correlation plot depicting changes in gene expression between all the elicitors evaluated in the comparative analysis: all transcriptomes compared at 30 min post treatment.

The observed transcriptional reprogramming induced by the COM treatment was compared to that induced (*i.e.* upregulated) by seven other well-characterized elicitors of plant immunity, recently reported by Bjornson *et al*. (Bjornson et al., 2021). The COM effect presents similarity with that of the other elicitors: 140 induced genes (∼30%) responded to COM and the other PTI elicitors (Fig. 4C). The transcriptional reprogramming induced by COM has however some uniqueness since 105 induced genes (∼20%) were not induced by any of the other tested elicitors (Fig. 4C). In fact, such level of specificity in transcriptional reprogramming was previously only observed for flg22 (Bjornson et al., 2021) (Fig. 4C). The lower number of induced genes by COM treatment could be related to the single time point used, differently from the other elicitors of PTI where multiple timepoints were used. The genes induced only by COM (and not by the other elicitors) were compared with genes found to be upregulated under abiotic stress (seven types of stresses were considered, *see Materials and Methods*). We observed that among these, 32 genes were induced solely by COM and not by any of the abiotic stresses (Supplemental Table S2 and Table S3); further suggestive of a certain degree of uniqueness on the COM’s effect.

The uniqueness of the COM treatment was further demonstrated through a correlation analysis of all elicitor transcriptomic datasets at the 30-min timepoint (Fig. 4D). At this timepoint, COM effect is not well correlated with any of the other tested elicitors; for example, bacterial hydroxy-fatty acid (3-OH-FA) and fungal chitooctamer (CO8). It also showed no correlation with the effect of oligogalacturonides (OGs) originating from plant cell wall pectin degradation. Cutin anchoring to the cell wall is a long-standing debate, but the involvement of polysaccharide-based moieties has been suggested (Philippe et al., 2020b). Polysaccharides can be found at very low amounts in cutin isolated using the ionic liquid extractant (Bento et al., 2021), but it remains an unresolved question if the detected polysaccharide-moieties are covalently linked to cutin. No glycoside-type linkages were detected in the NMR spectral fingerprint of a highly concentrated COM sample: 40 mg to allow the detection of low-intensity signals (Supplemental Fig. S5A). Several molecules derived from cell wall polysaccharides can act as elicitors, for example OGs (Ferrari et al., 2013), cellobiose (de Azevedo Souza et al., 2017), arabinoxylan oligosaccharides (Mélida et al., 2020) and mixed-linked β-1,3/1,4-glucans (Rebaque et al., 2021). However, the reported amounts for their eliciting effects (de Azevedo Souza et al., 2017) usually range from µg·mL^-1^ to mg·mL^-1^. These levels are higher than those detected in the COM preparations that were observed to contain only picograms of hydrolysable sugars *per* mg of COM (Supplemental Fig. S5B). The acquired data thus indicate that the molecules within the COM preparation acting as elicitors have a lipidic nature.

### Guiding principles on the chemistry of COMs that are able to elicit a rapid immune response in Arabidopsis

COM preparations have been shown to consist of oligomers and monomers (Escórcio et al., 2022). During infection, pathogens can secrete enzymes able to hydrolyze ester-type linkages present in cutin (Serrano et al., 2014); breaking the structural integrity of the cutin barrier to allow pathogen invasion of the infected plant tissue. To mimic such progressive attack of cutin, after obtaining a COM preparation, the non-hydrolyzed cutin fraction was recovered. The recovered cutin was subjected to a second round of mild hydrolysis and subsequently processed to obtain a COM II fraction. In *Arabidopsis* plants, the signal-intensity of the COM II induced calcium influx was >2-fold higher than that observed after COM treatment (Supplemental Fig. S6). This observation suggests that COM II might be more enriched in active elicitors compared to COM.

The free monomers were detected (and quantified) by GC-MS analysis, which also differentiates the methylated derivatives formed during the cleavage of esters through the methanolysis reaction (Supplemental, Table S1). The presence of oligomers was directly inferred from the detection of both primary (PAE) and secondary aliphatic esters (SAE) through NMR analyses, specifically in the HSQC spectral fingerprints of either COM preparation (Fig. 5A). The integration of their corresponding ^1^H NMR signals, relative to an internal standard, was used to infer the relative amount of PAE (*i.e.* linear) and SAE (*i.e.* branched) (Fig. 5B). Both types of esterification have been reported before in the spectral fingerprint of several cutin variants (Moreira et al., 2020; Bento et al., 2021; Escórcio et al., 2022). The estimated relative abundances of methyl esters in COM range from 40 to 70% of the total esters, consistent with the GC- MS data (Supplemental, Table S1). To depolymerize all present oligomers, the COM was subjected to hydrolysis and reanalyzed by GC-MS. Comparison of the resultant monomeric profiles of the COM and the resulting hydrolysate, exposed monomers increasing in abundance after hydrolysis (Supplemental, Table S1). The major aliphatic monomer of cutin, dihydroxy-C16 acid, is likely the major building block of the oligomers, distantly followed by 9,10-epoxy-18- hydroxyoctadecanoic acid, nonanedioic acid and hexadecanedioic acid.

**Fig. 5.**
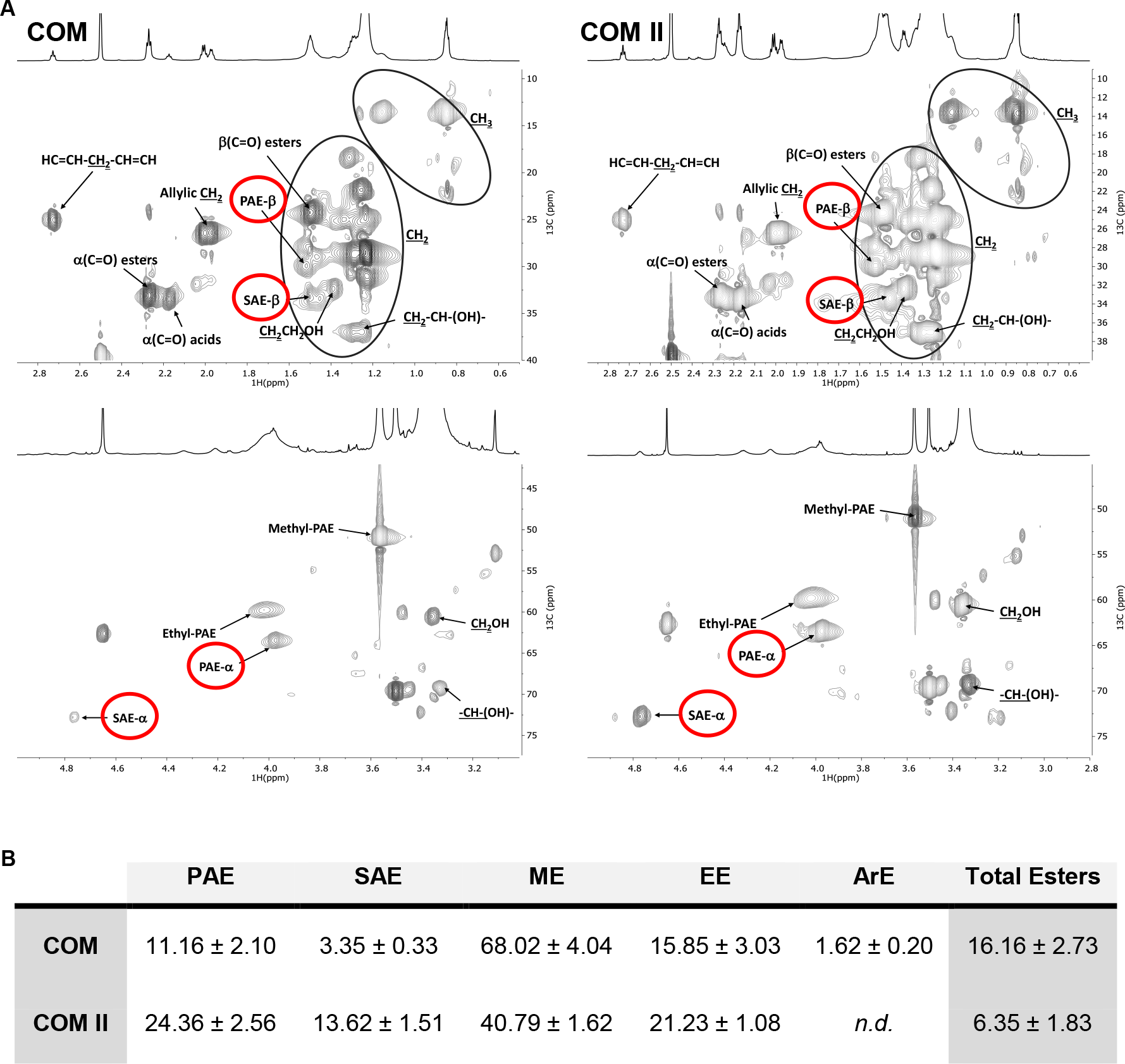
(A) NMR characterization of cutin-derived COMs. Magnification of the HSQC spectral regions corresponding to aliphatics for each sample. Some correlations (unlabelled) are uncertain or unidentified. **(B)** NMR quantification of the relative abundances of esters types present in each COM calculated through the integration of signals in the corresponding ^1^H NMR spectra. Ester types detected on this analysis include, PAE (Primary aliphatic esters), SAE (Secondary aliphatic esters), ME (Methyl esters), EE (Ethyl esters) and ArE (Aromatic Esters). Esters that were not detected on a sample are labelled as *n.d*.

A preliminary LC-MS/MS analysis was performed targeting the exact masses of dihydroxy-C16 acid dimers and trimers, carrying or not one methylation (Supplemental, Table S4A). Pure HPA was used to setup the method (see *Materials and Methods*). The given outputs (Compound Discovery 3.2) were unsupervised since the software automatically computes the most likely ions/adducts to be generated in negative/positive modes for each given mass (Supplemental, Table S4B). In both COMs, dimers were putatively identified, namely two dihydroxy-C16 acid molecules esterified, methylated or not – DP2 (Fig. 6A-B, Supplemental Fig. S7A-B). A trimer of dihydroxy-C16 acid molecules, carrying or not one methylation, was putatively identified only in COM II – DP3 (Fig. 6C-D, Supplemental Fig. S7C-D). These molecules can be a linear chain, yet one side-branch is possible (Fig. 6E-F). The NMR quantification data suggest that linear esters are in average two- to three-fold more abundant than branched esters (Fig. 5B), accordingly the linear DP3 chain is more likely to exist.

**Fig. 6.**
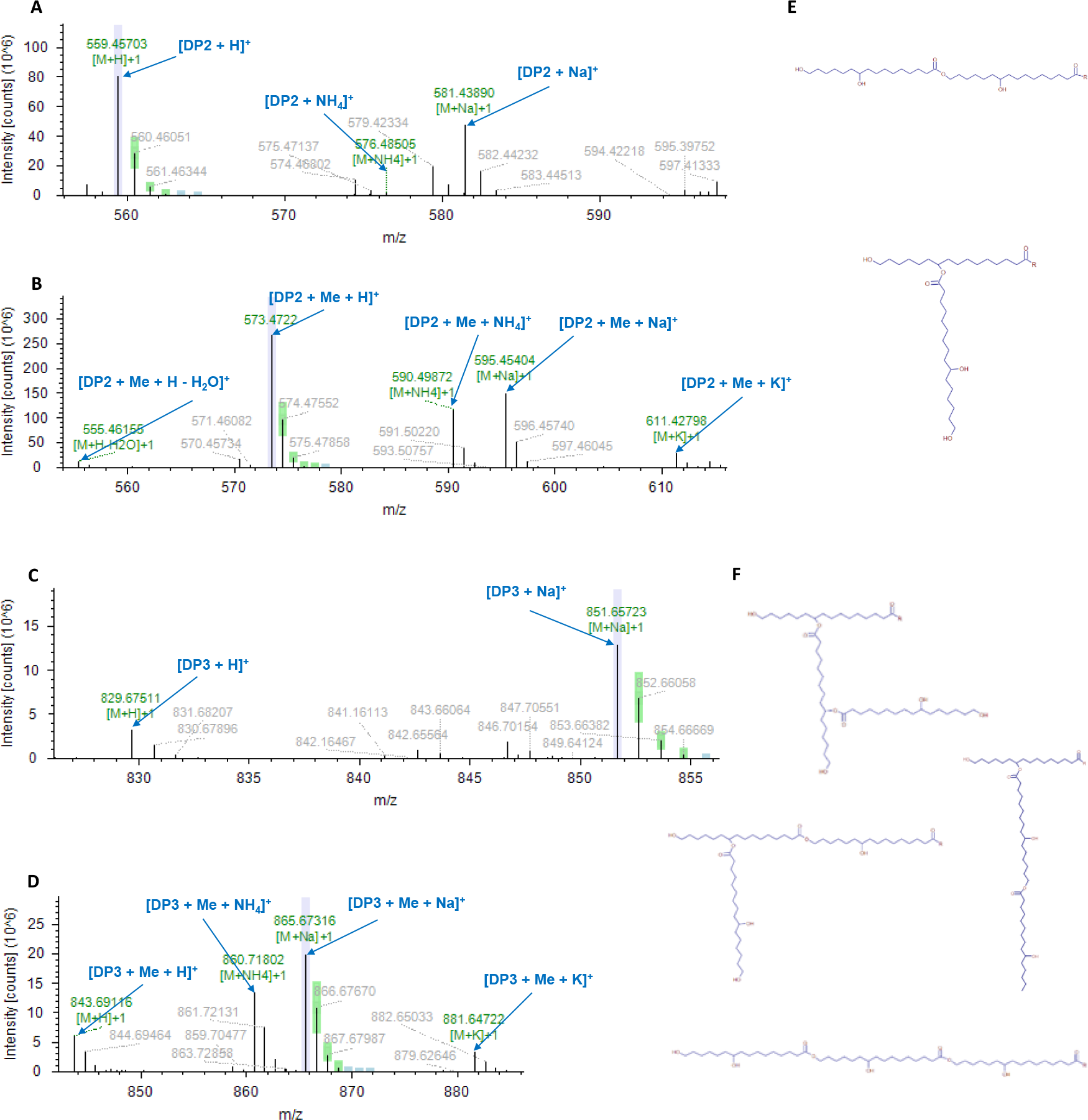
LC-MS/MS characterization of COM II (representative spectra for both COMs) in positive mode. MS1 spectra of the detected dimer composed by ester linked molecules of 10,16-dihydroxyhexadecanoic acid (DP2) in the non-methylated (**A**) and methylated (**B**) forms. MS1 spectra of the detected trimer composed by ester linked molecules of 10,16-dihydroxyhexadecanoic acid (DP3) in the non-methylated (**C**) and methylated (**D**) forms. Possible chemical configurations of the putatively identified oligomers (**E- F**), where R corresponds to an OH or CH3 group for the non-methylated and methylated forms, respectively. Lavender is presented when the labelled centroid matches the monoisotopic mass of the expected compound ion; Green is presented when the labelled centroid matches the delta mass and the relative intensity of the theoretic isotope pattern within the specified tolerances; Blue is presented when the expected centroid for this *m/z* value might be missing because its theoretic intensity is at the level of the baseline noise. The corresponding MS2 spectra are shown in Supplemental Fig. S7.

A MALDI-TOF method was developed to screen for the putative presence of oligomers up to octamers species (Supplemental Fig. S8). The MALDI-TOF analyses showed the presence of the most abundant free monomers in COM and COM II, some of which in the methylated form (Supplemental, Table S5). The controls - cutin and COMs hydrolysates - also contain the same non-methylated monomers (Supplemental, Table S5). In the COMs, dimers were identified, namely DP2, methylated or not (Fig. 7A), consistent with the LC-MS/MS. To selectively observe species capable of self-ionization such as aromatics, samples analyzed by MALDI-TOF were also analyzed without addition of ionization matrix (LDI-TOF). On the LDI-TOF spectra, other dimers detected consist of a dihydroxy-C16 acid esterified to coumaric acid without, or with one or two methylations - DP2c (Supplemental Fig. S9). The DP2c methylated molecules were only detected in the COM. The cutin hydrolysate (control) showed the presence of the non-methylated forms of DP2 in MALDI-TOF and DP2c in LDI-TOF (Supplemental, Table S5). NMR analyses of 40 mg of either COM, showed the presence of esterified aromatics only in COM (Supplemental, Fig. S10). Finally, the methylated-DP3 and, its non-methylated form, were identified in COM II (Fig. 7B), regardless of undetected in COM possibly due to lower relative abundance. Collectively the data on the COMs (and cutin hydrolysates) suggest that amongst the identified oligomeric species, the best PTI elicitor candidates are linear dimers or trimers composed of dihydroxy-C16 acid units, one of which possibly methylated.

**Fig. 7.**
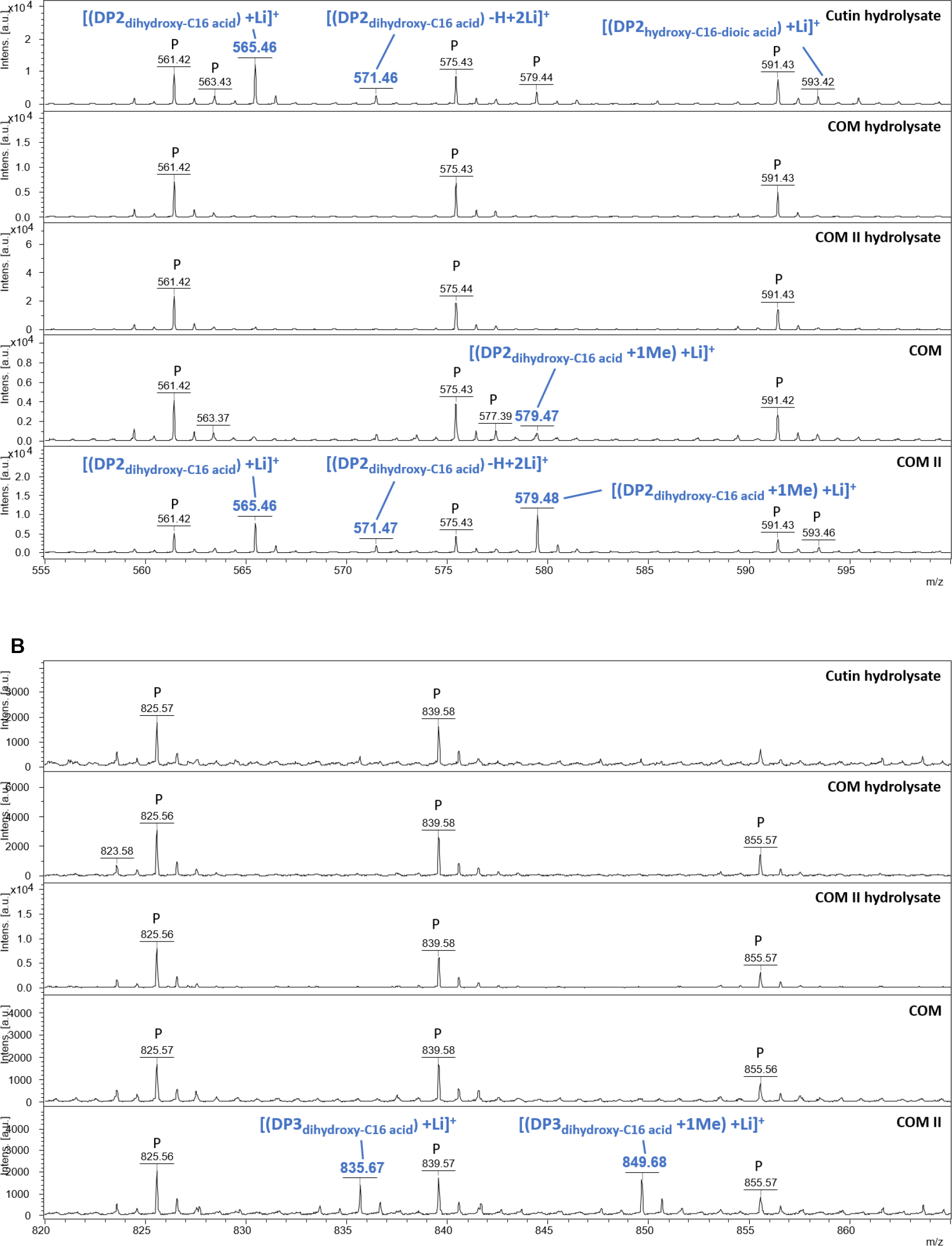
MALDI-TOF (+) spectra of cutin hydrolysate, COMs hydrolysates and COMs. The range 555-600 Da corresponds to expected masses for DP2s (**A**). The range 820-865 Da corresponds to expected masses for DP3s (**B**). Annotations were deduced from exact mass measurements. Black star indicates an ion from the MALDI matrix. “p” indicates an ion from PEG contamination.

## Conclusions

The plant cell wall barrier is an important interface during plant–microbe interactions, where cutin is the outermost polymeric component. Plants are able to recognize damages caused by pathogens, and elicit immune responses for example upon recognition of cell wall-derived fragments acting as DAMPs. As such, the cell wall barrier orchestrates key responses of the plant interaction with the surrounding environment. This rationale has defined the major hypothesis of our study, namely that cutin oligomers act as DAMPs able to trigger plant immune responses. In *Arabidopsis*, cutin oligomers (COMs), obtained through methanolysis of tomato pomace cutin, elicited several hallmark immune responses, including calcium influx (Fig. 2) and MAPK activation (Fig. 3), and a transcriptional response comprising features similar to those activated by well-characterized elicitors (Fig. 4). The perception mechanism of the COMs, which was observed to be independent of BAK1/BKK1 and CERK1 co-receptors, remains yet unresolved.

Chemical analyses identified that the COMs contain trimers and dimers (Fig. 6-7). The strongest elicitor candidates are the dihydroxy-C16 acid dimers (DP2) or trimers (DP3) carrying a methylation. DP2, methylated or not, were detected in both COMs. On the contrary, the DP3 and DP2c (dihydroxy-C16 acid esterified to coumaric acid), carrying or not methylation, were only detected in COM II and COM, respectively. The non-methylated DP2 and DP2c were present in cutin hydrolysates unable to elicit a calcium burst, though the threshold for PTI activation remains unknown. The methylation increases the oligomer’s lipophilicity, possibly favoring its diffusion; a hypothesis that requires focused analysis. Methyl-esters are, for example, present in seeds (Annarao et al., 2008) and vegetable oil (di Pietro et al., 2020). However, the isolated cutin polymer, which is deprived of methyl-esters, elicited a rapid ROS burst (Fig. 1A- B) and calcium influx (Fig. 2A). This observation questions the requirement of methylation for immune activation. Fungal lipases can generate methyl esters, for example from vegetable oil (Li et al., 2007). In plants, the modification of cutin degradation products by microbial methyl-transferases remains unknown in the context of PTI. However, methylation to potentiate the eliciting effect of cutin oligomers, may inspire alternative biotechnological valorization paths for fruit pomaces.

Both the cutin polymer (with minor degree of structural damage) and the generated oligomers acted as PTI elicitors. Previous work by others showed that some cutin monomers also activated some aspects of plant immunity (Kauss et al., 1999; Kim et al., 2008; Park et al., 2008). A step-by-step activation of specific elements of plant immunity by cutin having distinct degrees of structural damage, constitutes an appealing concept that deserves further investigation. The release of oligomeric elicitors during plant infection requires validation to attain a mechanistic understanding of cutin’s multiple functions in plant-pathogen interactions. The identity of the precise COM elicitors remain putative, and efficient syntheses are needed to obtain pure compounds. However, COMs clearly constitute a new class of DAMPs. Hence, their production from agro-industrial residues constitutes a promising value chain and may support development of sustainable agricultural bio-based treatments to increase disease resistance.

## Materials and Methods

### Plant Growth Conditions

*Arabidopsis thaliana* Col-0 seeds were germinated on soil and plants were grown for four-weeks in an Aralab Fitoclima climate chamber with 150 μmol·s^-1^·m^-2^ light intensity, following a 10 h/14 h day/night cycle, under constant temperature of 20 °C and 60 % humidity. Plants were watered automatically for 10 min three times *per* week. *Arabidopsis thaliana* Col-0 and the mutants *bak1-5 bkk1*, *cerk1-2^AEQ^* and *bbc* (Ranf et al., 2011; Roux et al., 2011; Xin et al., 2016), all in the Col-0 background, seeds were germinated on plates with 0.5 x Murashige and Skoog (MS) basal salt mixture supplemented with 1 % (w/v) sucrose and 0.9 % (w/v) phytoagar. After four days, seedlings were transferred to 24-well sterile culture plates containing 0.5x MS mixture supplemented with 1 % (w/v) sucrose and grown in sterile conditions in a Aralab Fitoclima climate chamber with 120 μmol·s^-1^·m^-2^ light intensity, following a 16 h/8 h day/night cycle, under temperatures of 20 °C during the day and 18 °C at night. The growth period was 8-days, 11-days or 14-days depending of the subsequent assays.

*Solanum lycopersicum* ‘Moneymaker’ seeds were germinated on soil and plants were grown for four weeks on a Conviron climate chamber with 120 μmol·s^-1^·m^-2^ light intensity, following a 12 h/12 h day/night cycle, under temperatures of 21 °C during the day and 19 °C at night and constant humidity of 60 %. Plants were watered manually three times *per* week.

*Nicotiana benthamiana* plants seeds were germinated on soil and plants grown for four weeks on a greenhouse room with 150 μmol·s^-1^·m^-2^ light intensity, following 16 h/8 h day/night cycle, under constant temperature of 24 °C. These plants were watered automatically daily for 20 min.

### Cutin Extraction

Cutin was extracted from tomato pomace as previously described(Moreira et al., 2020). The tomato pomace was obtained from Sumol + Compal, SA., and dried at 60 °C for one week until constant weight. Dry pomace was then milled using a Retsch ZM200 electric grinder (granulometry 0.5 mm; 10000 rpm) and stored at room temperature until further use. In brief, tomato pomace and cholinium hexanoate were mixed (1:10) and incubated for 2 h at 100 °C. The reaction was stopped by the addition of 80 mL of DMSO *per* gram of tomato pomace. The polyester was recovered by filtration using a nylon membrane filter (0.45 µm). Purification was obtained by washing with an excess of deionized water to remove all traces of DMSO. Purified cutin were then freeze dried and stored at room temperature for further use. Suspensions of the purified cutin powder were prepared in MiliQ water for testing purposes.

### Cutin Hydrolysis

To obtain a cutin oligomeric mixture (COM), a sodium methoxide-catalyzed methanolysis was performed by mixing 0.5 g of cutin with 20 mL of a solution of sodium methoxide(0.1M) in anhydrous methanol, at 40 °C for 2 h without stirring. At the end of the reaction, the mixture was cooled to room temperature and centrifuged (4 °C, 30 min, 4000 *g*) to recover the non- hydrolyzed cutin fraction. The supernatant (hydrolyzed fraction) was acidified to pH 3–3.5 by addition of HCl 37 % and subsequently centrifuged (4 °C, 30 min, 4000 *g*). The resulting precipitate was recovered, and the supernatant extracted three times by dichloromethane/water partition to release the hydrolysates; and sodium sulphate anhydrous was added to remove traces of water. The solution was concentrated under a constant nitrogen flux at 40 °C. To obtain cutin or COM hydrolysates, a sodium hydroxide alkaline hydrolysis was performed by mixing a solution of 0.5 M NaOH in methanol/water (1:1, v/v) at 95 °C with the cutin/COM powder, for 4 h. At the end of the reaction, the mixture was cooled to room temperature, then acidified to pH 3/3.5 with 1 M HCl, and subsequently extracted by dichloromethane/water partition to release the hydrolysable constituents. The solution was concentrated under a constant nitrogen flux at 40 °C.

### Immune assays

Leaf discs (collected using a 4-mm biopsy punch) or seedlings were transferred to white 96-well plates (one leaf disc or seedling *per* well) and equilibrated overnight in sterile ultrapure water (ROS measurements) or coelenterazine solution (Calcium measurements). The following day, the equilibration solution was removed, and replaced with a solution containing 100 µg.mL^-1^ up to 2 mg.mL^-1^ of COM, cutin hydrolysate, hydroxy palmitic acid (HPA) or cutin (Calcium measurements), and mixed with 1 mM luminol, and 10 µg.mL^-1^ HRP in the case of ROS measurements. Positive controls were also prepared with 100 nM (flg22 and Pep1) in MiliQ water, as well as blanks with 0.5 % (v/v) DMSO in MiliQ water or MiliQ water. Luminescence was detected and measured for 30 - 45 min using a Photek system equipped with a photon counting camera (leaf discs) or a Tecan Spark microplate reader (seedlings).

*MAPK activation* – For MAPK activation assays 14-day-old seedlings were used. The growth media was removed by inverting the plate on clean paper towels. Seedlings were treated for 10 min with 1 mL of 3 mg·mL^-1^ of COM, 100 nM flg22 (positive control) or the corresponding mock solutions (solvent control). Two seedlings *per* treatment were dried on clean paper towels, subsequently transferred to 1.5-mL tubes and instantly frozen in liquid nitrogen. All treated seedlings were stored at -80 °C until further use. Frozen seedlings were pulverized using a nitrogen-cooled plastic micro pestle, then mixed with 150 µL of extraction buffer containing 50 mM Tris-HCl pH 7.5, 150 mM NaCl, 2 mM EDTA, 10 %(v/v) glycerol, 2 mM DTT, 1 %(v/v) Igepal, and supplemented with protease and phosphatase inhibitors (equivalent to Sigma-Aldrich plant protease inhibitor cocktail and phosphatase inhibitor cocktails #2 and #3) was added. The tissue was then ground at 1800 rpm using an automatic stirrer fitted with a plastic micro pestle. The tubes were centrifuged at 15,000 *g* for 20 min at 4 °C in a refrigerated microcentrifuge. After centrifugation, 50 μL of extract were transferred to a fresh 1.5-mL Eppendorf tube. Samples were prepared for SDS-PAGE by heating at 80 °C for 10 min in the presence of 6x SDS loading buffer and 100 mM DTT.

Proteins were loaded to a 12 % (v/v) polyacrylamide gels, separated at 120 V for ≅ 120 min and subsequently transferred to a PVDF membrane at 100 V for 90 min at 4 °C. Membranes were then blocked for 2 h at room temperature or overnight at 4 °C in 5 % (w/v) milk in Tris buffered saline (50 mM Tris-HCl pH 7.4, 150 mM NaCl; TBS) containing 0.1 %(v/v) Tween-20 (TBS-T). Blots were probed in a 1:4000 dilution of the NEB anti-p42/p44-erk primary antibody in 5 % BSA in TBS-T for 2 h, followed by washing 4 times for 10 min each in TBS-T. Blots were then probed with a 1:10000 dilution of anti-rabbit secondary antibody in 5 % milk in TBS-T for 1 h, followed by washing 3 times for 5 min each in TBS-T. Finally, blots were washed for 5 min in TBS and treated with either standard ECL substrate or SuperSignal West Femto high sensitivity substrate (ThermoFisher Scientific). Blots were imaged using a Bio-Rad ChemiDoc Imaging System (Bio-Rad Laboratories).

### RNA Extraction and Sequencing

For RNA-seq experiments, 14-day-old seedlings were grown as described above. After nine days of growth in liquid MS medium supplemented with 1 % sucrose, the medium was removed from the wells and replaced with 600 µL of fresh liquid MS per well. The following day, 400 µl of 3 mg/mL of COM in 0.5 % DMSO in MiliQ water or the corresponding mock solution were added to each well. Seedlings were treated for 30 min and then two were collected and instantly frozen in liquid nitrogen. In total, four biological replicates were generated for each treatment (COM and mock) and stored at -80 °C for further processing.

Frozen seedlings were pulverized while frozen using a Spex SamplePrep Geno Grinder 2010 at 1500 rpm for 90 s. Total RNA was extracted at 4 °C from two ground seedlings as previously described(Shi and Bressan, 2006) by addition of 900 µL of TRI reagent (Ambion) and 200 µL of chloroform, recovery of 400 µL from the aqueous phase, precipitation with 500 µL of isopropanol and washing with 70 % ethanol. All samples were then solubilized in 30 µL of RNase-free water. Samples were subsequently subjected to DNase treatment using a TURBO DNA-free Kit (Ambion) according to manufacturer’s instructions. The reaction mix was incubated at 37 °C for 30 min, after which the inactivation reagent was added and incubated for 5 min at room temperature. After centrifugation the supernatant was transferred to a new tube. Quantification and quality assessment of all RNA samples were evaluated on a TapeStation (Agilent) and RNA sequencing performed by the Beijing Genomics Institute (BGI).

### RNA-seq data processing

For paired-end RNA sequencing (RNA-seq), libraries were generated at BGI according to the DNBSEQ stranded mRNA library system. Eight samples were indexed and sequenced using the DNBseq™ sequencing platform (20 million reads per sample). Generated FastQ files were analyzed with FastQC, and any low-quality reads were trimmed with Trimmomatic (Bolger et al., 2014).

All libraries were aligned to the *A. thaliana* genome assembly TAIR10 with gene annotations from Ensembl Plants v.49 using the HISAT2 v.2.1.0 pipeline(Kim et al., 2015) followed by read counts with HTSeq v. 0.13.5(Anders et al., 2015). All RNA-seq experiments were carried out with four biological replicates. Differential expression analysis, and quality control principal- component analysis (PCA) and MA plots were generated using the DESeq2 v.1.30.0 R package(Love et al., 2014). The genes that showed |log2| > 1-fold changes in expression with an adjusted P value below 0.05 are defined as significantly differentially expressed genes (DEGs) in this analysis. Transcript abundance was defined as transcripts per kilobase million (TPM). Gene Ontology enrichment of the differentially expressed genes was performed with the topGO v.2.42.0 R package, using the Fisher exact test to attain significantly enriched categories.

### Comparative analysis of transcriptome modification upon elicitor treatment

Differentially expressed gene lists in response to seven elicitors (3- OH-FA, CO8, elf18, flg22, nlp20, OGs, Pep1) upon treatment under similar conditions as COM were obtained from Bjornson, *et al*. (2021) (Bjornson et al., 2021). This study followed a time course from 5 min to 3 h post-elicitation: a gene list was obtained for each elicitor with genes significantly induced at any time. Abiotic stress treatment analysis for seven abiotic stresses (heat, cold, drought, salt, high osmolarity, UV-B light, wounding) was also obtained from Bjornson, *et al*. (2021), based on ATH1 microarray experiments presented in Killian, *et al*. 2007 (Kilian et al., 2007). This study followed a time course from 5 min to 12 h post-elicitation: a gene list was obtained for each elicitor with genes significantly induced at any time up to 3 h. Comparisons and visualizations among differentially expressed genes were performed in R using the tools of the tidyverse v.1.3.1 package (Wickham et al., 2019). Spearman correlation among log_2_ (fold changes) for treatments was calculated using the Hmisc package in R v.4.5-0) and visualized using the corrplot package v.0.89. Annotation data for genes induced specifically by COM was obtained from the *Arabidopsis* information resource (TAIR) via Bioconductor package org.At.tair.db v.3.10.0. The raw data and the processed file are deposited in the ArrayExpress (www.ebi.ac.uk/fg/annotare/edit/16708/), publicly available after curation.

### Quantitative analyses of total carbohydrate content

To evaluate the polysaccharide content, each COM sample was subjected to an acid hydrolysis (1 M H_2_SO_4_ in methanol) for 4h at 90 °C. The hydrolysable sugars were recovered in the supernatant through centrifugation (18514 *g*, 4 °C, 20 minutes) and the pH was neutralized using 5 M NaOH in water. All samples were dried under a flux of nitrogen at room temperature. Quantification of carbohydrates in the dried hydrolysates was performed using the total carbohydrate assay kit from Sigma-Aldrich according to the manufacturer’s instructions. The samples were analyzed in triplicates.

### NMR characterization of cutin oligomeric mixtures

NMR spectra of COMs were recorded using an Avance III 800 CRYO (Bruker Biospin, Rheinstetten, Germany). All NMR spectra (^1^H, ^1^H-^1^H COSY, ^1^H-^13^C HSQC, ^1^H-^13^C HMBC) were acquired in DMSO-*d*_6_ using 5 mm-diameter NMR tubes, at 60 °C as follows: 15 mg of COMs in 400 μL of DMSO-*d*_6_ (in triplicate) or for validation 40 mg of COMs in 400 μL of DMSO-*d*_6_. For quantification purposes, 1.25 mg of benzene (internal standard) was added to each sample. MestReNova, Version 11.04-18998 (Mestrelab Research, S.L.) was used to process the raw data acquired in the Bruker spectrometers.

### GC-MS characterization of cutin oligomeric mixtures

To release the hydrolysable constituents, the COMs were treated with a solution of 0.5 M NaOH in methanol:water (1:1 [v/v]) at 95 °C for 4 h. The mixture was cooled to room temperature and acidified to pH 3–3.5 with HCl 1 M, spiked with a known concentration of hexadecane (internal standard), and extracted three times with dichloromethane. Sodium sulphate anhydrous was added to the organic phase to remove water and concentrated under a nitrogen flow. The non-hydrolysable fraction was recovered by filtration (cellulose nitrate filter) and subsequently washed, dried, and weighted (recalcitrance). The COMs samples were also analyzed directly, *i.e.* not subjected to alkaline hydrolysis. The dried samples were derivatized in N,O-bis(trimethylsilyl)trifluoroacetamide containing 1 % (v/v) of trimethylchlorosilane in pyridine (5:1), for 30 min at 90 °C. The derivatives were then analyzed by GC-MS (Agilent: 7820A GC and 5977B quadrupole MS; HP-5MS column) as follows: ramp temperature 80 °C, then 2 °C·min^-1^ to 310 °C for 15 min. The MS scan mode, with source at 230 °C and electron impact ionization (EI+, 70 eV) was used for all samples. Data acquisition was accomplished by MSD ChemStation (Agilent Technologies); compounds were identified based on the equipment spectral library (Wiley-National Institute of Standards and Technology) and quantified using external standards of the major classes of the aliphatic monomers (heptadecanoic acid, hexadecanedioic acid and pentadecanol). All samples were analyzed in triplicates, each with technical duplicates.

### LC-MS/MS characterization of COMs

The LC-MS/MS protocol was adapted from Bhunia R. *et al*. (2018) (Bhunia et al., 2018). The experiments were performed in a Q Exactive Focus™ Hybrid Quadrupole-Orbitrap™ Mass Spectrometer coupled to a Dionex Ultimate 3000 UHPLC. HPA was used as a standard and prepared in isopropanol:methanol:acetonitrile (1:1:1) at a concentration of 200 ng/µL. The samples were prepared in the same way at 1 µg/µL. Separation was achieved in a Waters XBridge column C18 (2.1x150 mm, 3.5 µm particle size, P/N 186003023), using a gradient of increasing percentage of 20 mM ammonium formate in isopropanol (IPA): acetonitrile (ACN) (75:25) (B) and decreasing percentage of ACN:water (60:40) with 20 mM ammonium formate (A). The total method time was 77 min, the flow rate was 0.4 mL·min^-1^, and the column was kept at 37 °C. The data was acquired using the Xcalibur software v.4.0.27.19 (Thermo Scientific). The method consisted of several cycles of Full MS scans (R=70000; Scan range=100-1500 m/z) followed by 3 ddMS2 scans (R=17500; NCE 30 V) in positive and negative mode. External calibration was performed using LTQ ESI Positive/Negative Ion Calibration Solution (Thermo Scientific). Generated mass spectra were processed using Compound Discoverer 3.2 (Thermo) for small molecule identification. The search was performed against the mass list with provided molecular formulas (dimers, trimers), as well as mzCloud MS2 database, KEGG and ChEBI MS1 databases. A 3-ppm mass tolerance was used. The minimum peak intensity (MS1) for detection was 10^6^. A manual validation of the assignments for the identified oligomers was performed by inspection the MS2 fragmentation profiles against the theoretical fragmentation generated on Mass Frontier 8.0 (Thermo). Theoretical chemical structures for the identified oligomers were generated ChemDraw 21.0.0.

### LDI-TOF and MALDI-TOF analyses of COMs and corresponding hydrolysates

The samples were analyzed by laser desorption/ionization (LDI)- time-of-flight (TOF) MS and by matrix-assisted laser desorption/ionization (MALDI)-time-of-flight (TOF) MS. As control for monomers, total alkaline hydrolysate of cutin was used. As oligomers control, a batch of oligomers (DP1 to DP8) were produced (Supplemental Fig. S8) from purified cutin monomers as previously described (Marc et al., 2021) with small modification. The polymerization time was reduced to 8 h and the oligomers were extracted from the polymer by hot (70 °C) ethanol extraction.

For the LDI-TOF analyses, the samples were deposited on a polished steel MALDI target plate and analyzed without any preparation. For the MALDI-TOF analyses, samples were mixed with the matrix solution composed of DHB (2,5-dihydroxybenzoic acid) 3 mg·mL^-1^ in 75% methanol, with 2.5 mM LiCl, in a 1:3 ratio (v/v). The mixture (1 μL) was deposited on a polished steel MALDI target plate. Measurements were performed on a rapifleX MALDI-TOF spectrometer (Bruker Daltonics, Bremen, Germany) equipped with a Smartbeam 3D laser (355 nm, 10000 Hz) and controlled using the Flex Control 4.0 software package. The mass spectrometer was operated in reflectron mode with positive polarity for MALDI-TOF analyses and in negative polarity for LDI- TOF analyses. Spectra were acquired in the range of 180-5000 m/z. Neither the MALDI-TOF nor the LDI-TOF used in these experiments can observe the signal of the free *p*-coumaric acid.

## Acknowledgements

The authors are thankful to Pedro Lamosa (ITQB NOVA) for support with the NMR analyses. Mass spectrometry data were generated by the Mass Spectrometry Unit (UniMS), ITQB/iBET, Oeiras, Portugal. We thank Stefanie Ranf for the gift of *cerk1-2^AEQ^*seeds. All members of the Silva Pereira and Zipfel labs are also thanked for useful discussions.

## Authors’ contributions

CSP and CZ supervised the project and the interpretation of data; CSP prepared the final version of the manuscript. All authors have made substantial contributions to the acquisition, analysis and interpretation of data and contributed to the drafting of the manuscript: CJSM, AB and RE (cutin and COM preparation); CJSM (all plant experiments); CJSM, MB and CM (RNA seq); AST, CJSM and RE (GC-MS); AB and RE (NMR); IM and CJSM (LC-MS/MS); BB and MF (MALDI-TOF); CJSM (preparation of the initial draft of the manuscript). All authors read and approved the final version of the manuscript.

